# Investigating the Interplay Between Affective, Phonatory and Motoric Subsystems in Autism Spectrum Disorder Using a Multimodal Dialogue Agent

**DOI:** 10.1101/2021.04.10.439293

**Authors:** Hardik Kothare, Vikram Ramanarayanan, Oliver Roesler, Michael Neumann, Jackson Liscombe, William Burke, Andrew Cornish, Doug Habberstad, Alaa Sakallah, Sara Markuson, Seemran Kansara, Afik Faerman, Yasmine Bensidi-Slimane, Laura Fry, Saige Portera, David Suendermann-Oeft, David Pautler, Carly Demopoulos

## Abstract

We explore the utility of an on-demand multimodal conversational platform in extracting speech and facial metrics in children with Autism Spectrum Disorder (ASD). We investigate the extent to which these metrics correlate with objective clinical measures, particularly as they pertain to the interplay between the affective, phonatory and motoric subsystems. 22 participants diagnosed with ASD engaged with a virtual agent in conversational affect production tasks designed to elicit facial and vocal affect. We found significant correlations between vocal pitch and loudness extracted by our platform during these tasks and accuracy in recognition of facial and vocal affect, assessed via the Diagnostic Analysis of Nonverbal Accuracy-2 (DANVA-2) neuropsychological task. We also found significant correlations between jaw kinematic metrics extracted using our platform and motor speed of the dominant hand assessed via a standardised neuropsychological finger tapping task. These findings offer preliminary evidence for the usefulness of these audiovisual analytic metrics and could help us better model the interplay between different physiological subsystems in individuals with ASD.

## 1. Introduction

Autism Spectrum Disorder (ASD) is a neurodevelopmental condition that manifests as deficits in social communication and interaction [1]. Epidemiological surveys estimate the prevalence of ASD in the United States at 18.5 per 1,000 children [2]. Autism diagnosis rates have increased due to an increase in awareness and access to resources. In the state of California, the diagnosed autism incidence rate increased by 612% in the past two decades [3]; however, sociodemographic barriers such as race, economic challenges and stigma still hamper early detection and intervention in children with autism [4].

Individuals with ASD are known to have impaired processing of emotions (affect recognition) and impaired production of both vocal and non-verbal emotional expressions (affect production) during communication [5, 6, 7]. Acoustic characteristics like prosody and fundamental frequency of speech in children with ASD are often perceived as atypical when compared to speech in typically-developing children [8, 9]. Individuals with ASD produce emotional phrases that are louder, longer, more variable in pitch, and sound less natural [10]. Facial affect production has been demonstrated to be equally difficult for individuals with ASD in both imitation and expression tasks [11]. Also, facial expressions of emotion are rated as more intense and less natural in individuals with ASD [12] by both neurotypical raters as well as raters with ASD [13].

Prior work has demonstrated the objective significance of automated quantitative assessment of atypical speech production and facial expression in ASD [14, 15, 16]. Such automated assessments can help clinicians objectively quantify skills in this domain of functioning for which no standardised, age-normed, validated measures are available. [17, 18, 19].

Moreover, the coordination of facial and vocal expressions during emotional speech production is less coordinated in ASD [20]. Due to such cross-domain atypicalities in ASD, a multi-modal framework approach has been suggested to provide useful quantitative insights in early diagnosis and categorisation of ASD [21, 22, 23]. In this paper, we explore the interplay between multiple neuropsychological subsystems through examination of automatically-extracted voice and facial metrics during a novel conversational task and a comprehensive set of clinically-validated objective measures. We probe the feasibility of a scalable, low-cost and remotely-administrable multi-modal conversational platform for this purpose. Such technology could potentially assist clinicians and researchers in gathering relevant diagnostic information and in longitudinally monitoring children with developmental disorders.

## 2. Multimodal Conversational Platform

NEMSI or NEurological and Mental health Screening Instrument is a cloud-based multimodal dialogue technology in which participants engage in a conversation with a virtual agent and speech and facial behaviours are elicited through a variety of exercises. Data analytics modules automatically extract relevant speech and facial metrics from the captured audiovisual data in real time and store them in a database. These metrics, with a detailed participant-wise, session-wise and task-wise breakdown, can be accessed by researchers via a user-friendly dashboard. A more detailed description of NEMSI, conversational protocols and a schematic of the software and hardware architecture of the system can be found in our previous publications [24, 25, 26].

## 3. Methods

The current study was conducted on site at the University of California, San Francisco and was approved by the universitys’ Institutional Review Board. Informed consent was obtained from the participants’ guardians along with a written assent from the participants. Data from 22 participants (10 females, mean age: 11.37 ± 2.47 years, see Figure 1) diagnosed with ASD by a licensed clinical psychologist was included in the analysis.

**Figure 1:**
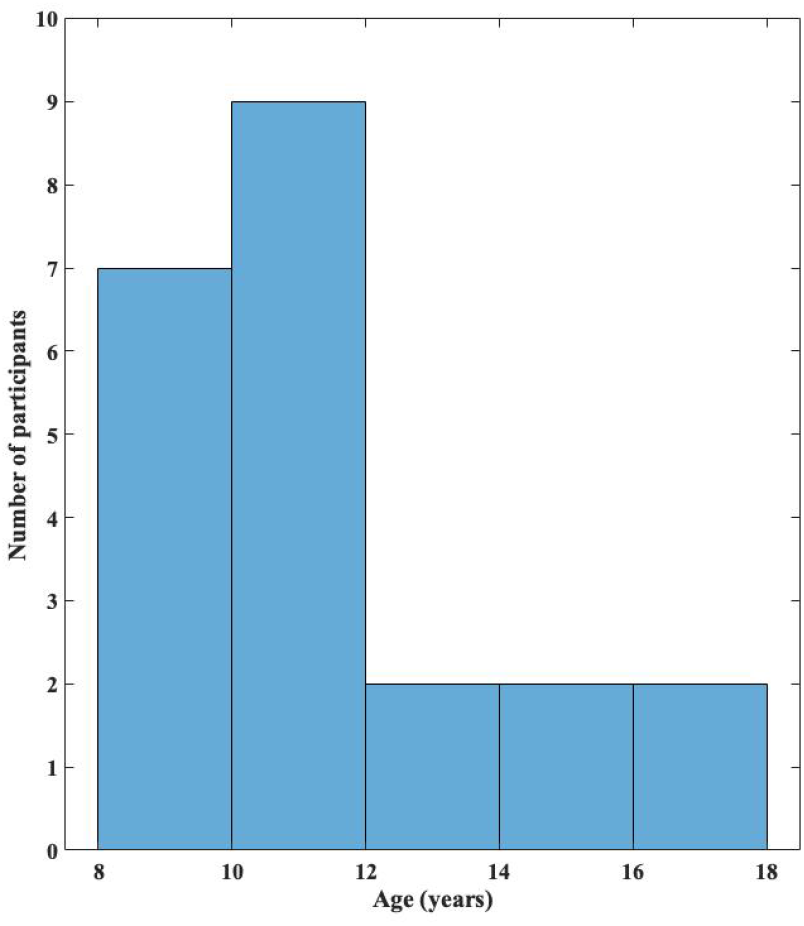
Distribution of age in the cohort

### 3.1. Standardised Clinical Instruments

All participants / guardians underwent a thorough neuropsychological evaluation battery including the Autism Diagnostic Observation Schedule, Second Edition (ADOS-2) [27], the Autism Diagnostic Interview - Revised [28], the Behavior Assessment System for Children, Third Edition (BASC-3) [29], the Diagnostic Analysis of Nonverbal Accuracy (DANVA) [30, 31] and NEPSY-II [32]. The ADOS-2 scores for 11 participants were collected during the COVID-19 pandemic and were deemed invalid because the participants wore a mask during the assessment. These scores were excluded from the analysis. Additionally, participants also took part in a motor skills and fine motor dexterity assessment, including bilateral manual grip strength assessed with a Lafayette Hand Dynamometer Model J00105 [33], finger tapping speed assessed with a standard board-mounted finger tapper with counter [34], and the Grooved Peg-board Test [35]. All scores were age scaled based on the referenced norms.

### 3.2. Affect Production Task

The Affect Production Task (APT) was administered via Modality.AI’s NEMSI platform. Participants were prompted to produce four emotions (happy, sad, angry and afraid) through five tasks:

1. Monosyllabic Emotion Production (8 turns, 2 turns per emotion): a video stimulus prompted the users to say “oh” in a way that conveyed the specified emotion during every turn;
2. Sentence-length Emotion Production (8 turns, 2 turns per emotion): a video stimulus prompted the users to say “I’ ll be right back” in a way that conveyed the specified emotion during every turn;
3. Emotion-eliciting Situations (16 turns, 4 turns per emotion): the virtual agent narrated a situation and a corresponding emotional response at the end of which a picture depicting the situation was shown and the user was prompted to say “oh” in a way that conveyed the specified emotion that was situationally appropriate;
4. Monosyllabic Repetition (16 turns, 4 turn per emotion): an audio recording of “oh” in one of the four emotions was played during every turn and the user was asked to repeat the monosyllabic production in the same emotional inflection;
5. Facial Repetition (8 turns, 2 turns per emotion): a video prompted the user to make a face representing one of the four emotions like the person in the video during every turn.

### 3.3. Analytics

Speech metrics (Fundamental Frequency in Hz, Articulation Loudness in dB) and facial metrics (Eyebrow Height in pixels, Eye Openness in pixels, Lip Aperture in pixels, Mouth Surface Area in pixels^2^, Jaw Velocity in pixels/frames and Jaw Acceleration pixels/frames^2^) were calculated in real time [25] and displayed on a dashboard accessible to the clinicians and the researchers involved in the study. Facial metrics in pixels were normalised within every subject by dividing the values by the inter-lachrymal distance in pixels for each subject.

### 3.4. Statistical Methods

Linear correlations were run between measures from standardised neuropsychological assessments and the speech and facial metrics derived from our platform. To control for false positives due to multiple comparison, the Benjamini-Hochberg procedure

[36] was used to adjust the p-values using a false discovery rate of *α* = 0.05. Findings that survived this adjusted significance threshold were included in this paper.

## 4. Observations

### 4.1. Language Level and IQ

The Clinical Evaluation of Language Fundamentals-Fifth Edition (CELF-5) [37] and the Weschler Intelligence Scale for Children - Full Scale IQ (WISC - FSIQ) [38] scores for the cohort are reported in Table 1.

**Table 1:** Language level and IQ measures in the cohort

| Measure | Range | Mean ± SD |
| --- | --- | --- |
| CELF-5 Core Language Score | 59 - 143 | 100.77 ± 22.06 |
| CELF-5 Expressive Language Index | 59 - 131 | 100.09 ± 20.48 |
| CELF-5 Receptive Language Index | 58 - 141 | 100.73 ± 23.08 |
| WISC - FSIQ | 50 - 136 | 102.95 ± 19.79 |

### 4.2. Speech Metrics

We found significant positive correlations between the DANVA Adult Receptive Paralanguage Subscore (APSS) and the fundamental frequency of the participants’ speech during the Emotion-Eliciting Situation subtask in the APT (Happy: r^2^ = 0.41 and Sad: r^2^ = 0.54) and also in the Monosyllabic Emotion Production subtask (Happy: r^2^ = 0.56 and Angry: r^2^ = 0.44). We also found positive correlations between the DANVA Child Receptive Paralanguage Subscore (CPSS) and the fundamental frequency of the participants’ speech during the Emotion-Eliciting Situation subtask (Happy: r^2^ = 0.55 and Sad: r^2^ = 0.45), the Monosyllabic Emotion Production subtask (Happy: r^2^ = 0.48) and the Monosyllabic Repetition subtask (Angry: r^2^ = 0.36). Fundamental frequency of speech during all these tasks did not correlate with participants’ age so the effect of age on these correlations can be ruled out. Additionally, the DANVA CPSS score was also positively correlated with Articulation Loudness during the Sentence-length Emotion Production subtask (Happy: r^2^ = 0.34). Please refer to Table 2 for more details.

**Table 2:**
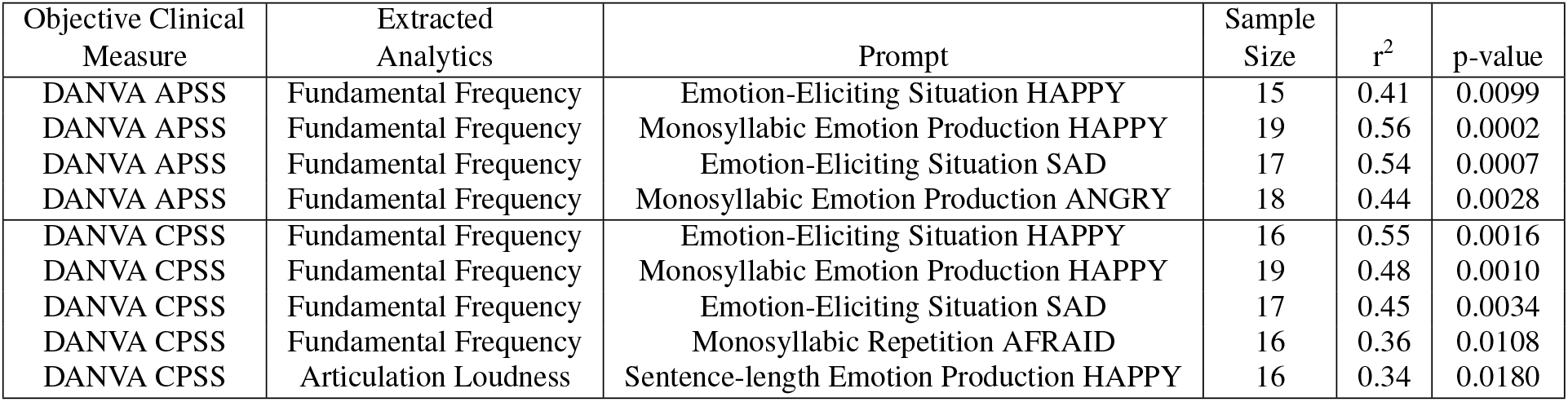
Speech acoustic measures of fundamental frequency (F0) and articulation loudness extracted automatically correlate with the participants’ accuracy at recognition of facial and vocal affect in audio recordings of adults and children

### 4.3. Facial Metrics

Motor speed of the dominant hand assessed using a finger tapping task was found to be positively correlated with participants’ maximum jaw velocity during the Monosyllabic Repetition subtask in the APT (Sad: r^2^ = 0.43 and Afraid: r^2^ = 0.37) and during the Emotion-Eliciting Situation subtask (Afraid: r^2^= 0.36). Motor speed of the dominant hand was also positively correlated with maximum jaw acceleration during the Emotion-Eliciting Situation subtask (Afraid: r^2^ = 0.42). Please refer to Table 3 for more details.

**Table 3:**
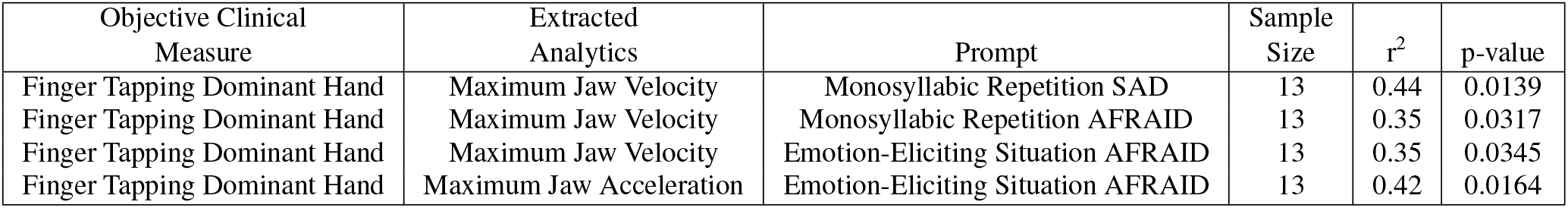
Jaw kinematic measures (velocity and acceleration) extracted automatically correlate with motor speed of the dominant hand

| Objective Clinical Measure | Extracted Analytics | Prompt | Sample Size | $r^2$ | p-value |
| --- | --- | --- | --- | --- | --- |
| Finger Tapping Dominant Hand | Maximum Jaw Velocity | Monosyllabic Repetition SAD | 13 | 0.44 | 0.0139 |
| Finger Tapping Dominant Hand | Maximum Jaw Velocity | Monosyllabic Repetition AFRAID | 13 | 0.35 | 0.0317 |
| Finger Tapping Dominant Hand | Maximum Jaw Velocity | Emotion-Eliciting Situation AFRAID | 13 | 0.35 | 0.0345 |
| Finger Tapping Dominant Hand | Maximum Jaw Acceleration | Emotion-Eliciting Situation AFRAID | 13 | 0.42 | 0.0164 |

## 5. Discussion

In this study, we found that speech metrics derived by NEMSI, a cloud-based multimodal dialogue technology, are significantly correlated with recognition of vocal affect assessed via the Diagnostic Analysis of Nonverbal Accuracy (DANVA) task. We also found correlations between jaw kinematics and motor speed of the dominant hand as assessed by a standardised finger tapping task.

The DANVA receptive paralanguage subtasks assess the participants’ ability to identify vocal emotional communication, with higher scores indicating competence in recognition of vocal affect. Participants who were more competent in identifying the emotion from adult and child voice recordings produced emotional speech (monosyllabic “oh”) with a higher fundamental frequency. A previous study has found an increased pitch range in high-functioning autism [9]. Participants with better DANVA CPSS scores also produced louder sentence-length happy emotional speech. Increased articulation loudness is indeed associated with happy emotional speech [39]. Our findings may suggest that high-performing participants on the vocal affect recognition task may possess greater ability to modulate the acoustic properties of emotional speech whereas more impaired participants are unable to do so. This may suggest that there is a relationship between the recognition and production of the acoustic properties that are relevant for signifying emotion. Indeed, previous research has identified a relationship between basic auditory processing and vocal affect recognition [40, 41, 42].

The correlations between jaw movement and motor dexterity of the dominant hand may be a manifestation of global motor impairments in individuals with ASD [43, 44]. Motor deficits in ASD have been attributed to abnormal sensorimotor integration [45, 46] due to increased sensory noise [47] or dysfunctional representation of internal models of action [48]. A previous study has found no differences in finger-tapping accuracy between children with ASD and typically-developing children but there was increased variability in temporal processing parameters in the ASD group [49]. Correlations between jaw movement and finger tapping suggest global altered temporal processing in ASD which may also explain poor cross-modal coordination during emotional speech production [20]. Indeed, prior work has suggested a coupling between speech motor co-ordination and fine motor skill systems in ASD [50].

Our preliminary results indicate that speech and facial metrics extracted from our APT task administered using Modality.AI’s NEMSI platform can tap into deficits of various physiological subsystems in children with ASD. The current study was conducted in a laboratory setting but the conversational platform can also be accessed remotely via a weblink [26]. Thus, our technology provides the opportunity for clinicians and researchers to monitor children with developmental disorders in a non-clinical, non-laboratory setting. Since deficits in emotion recognition have been observed cross-culturally in individuals with ASD [51], our technology can also be leveraged to compile a global corpus of speech and facial metrics during affect recognition and production in ASD. A conversational task where participants are prompted to produce a monosyllabic “oh” may also prove useful in a low-functioning population. Furthermore, it has been shown that children with ASD can be trained to produce qualitatively-improved emotional facial expressions [52]. Remote monitoring may also facilitate a longitudinal, quantitative analysis of such training paradigms.

Future studies would involve collecting data from larger cohorts of children with ASD along with data from typically-developing children to examine whether differences in vocal and facial affect production between the two groups can be captured through automatically-extracted metrics.

## 6. Acknowledgements

This study was supported by the National Institutes of Health grant K23 DC016637 and Autism Speaks grant 11637 awarded to Carly Demopoulos.

